# Novel modification of Luminex assay for characterization of extracellular vesicle populations in biofluids

**DOI:** 10.1101/2022.01.12.475897

**Authors:** OV Volpert, E Gershun, K Elgart, V Kalia, H Wu, AA Baccarelli, E Eren, D Kapogiannis, A Verma, A Levin, E Eitan

## Abstract

Most approaches to extracellular vesicle (EV) characterization focus on EV size or density. However, such approaches provide few clues regarding EV origin, molecular composition, and function. New methods to characterize the EV surface proteins may aid our understanding of their origin, physiological roles, and biomarker potential. Recently developed immunoassays for intact EVs based on ELISA, NanoView, SIMOA and MesoScale platforms are highly sensitive, but have limited multiplexing capabilities, whereas MACSPlex FACS enables the detection of multiple EV surface proteins, but requires significant quantities of purified EVs, which limits its adoption. Here, we describe a novel Luminex-based immunoassay, which combines multiplexing capabilities with high sensitivity and, importantly, bypasses the enrichment and purification steps that require larger sample volumes. We demonstrate the method’s specificity for detecting EV surface proteins using multiple EV depletion techniques, EVs of specific cellular origin isolated from culture media, and by co-localization with established EV surface markers. Using this novel approach, we elucidate differences in the tetraspanin profiles of the EVs carrying erythrocyte and neuron markers. Using size exclusion chromatography, we show that plasma EVs of putative neuronal and tissue macrophage origin are eluted in fractions distinct from those derived from erythrocytes, or from their respective cultured cells. In conclusion, our novel multiplexed assay differentiates between EVs from erythrocytes, macrophages, and neurons, and offers a new means for capture, classification, and profiling of EVs from diverse sources.

## Introduction

Extracellular vesicles (EVs) are released by all cells in the body via diverse mechanisms including outward budding of the plasma membrane/exocytosis (microvesicles) and reverse-inward budding of endosomal membranes (exosomes) (Rak 2010; Colombo et al. 2014). Regardless of their biogenesis pathway, it is widely accepted that plasma and other biological fluids contain EVs arising from multiple cell types. Furthermore, a large body of evidence shows that EV molecular composition closely reflects that of the cell of origin (He et al. 2018). Importantly, EV surface proteins also reflect their cell(s) of origin, and potentially also biogenesis pathways, and determine EV destination, and function (Colombo et al. 2014).

The development of flow cytometry for the detection of surface proteins on immune cells represented a landmark in immune and cellular biology. The development of a similar approach that extends this capacity towards EV analysis may prove equally important. To address this experimental and knowledge gap, a number of assays were developed to characterize intact EVs (Kanwar et al. 2014; Ohmichi et al. 2019; Ter-Ovanesyan et al. 2021). All these assays are based on a sandwich immunoassay principle, wherein the sandwich is formed by an EV and two antibodies targeting distinct proteins on the EV surface. To generate a signal, both antigens should be interconnected, presumably via the EV membrane. This principle can be incorporated into multiple immunoassay platforms including traditional Enzyme-Linked Immunosorbent Assay (ELISA), Meso Scale Discovery (MSD) electrochemiluminescence, Single Molecule Array (SIMOA), flow cytometry, Surface Plasmon Resonance (SPR) and many others. However, many of these methods provide limited capabilities of simultaneous examination of multiple EV surface proteins (Nolan and Duggan 2018; Chiang and Chen 2019; Kurian et al. 2021). The exception is the MACSPlex FACS Exosome Kit (Miltenyi Biotech, cat. No 130-108-813), which enables the detection of multiple EV surface epitopes (a total of 37). However, its capacity for simultaneous assessment is limited by the number of compatible fluorescent dyes and dye interference. In addition, this is an arduous methodology that requires EV enrichment prior to analysis and highly proficient and skilled operators.

Here, we describe the first intact EV immunoassay based on the Luminex platform (Fig. 1A). Our technique, intact EV Luminex (IEL) utilizes MagPlex spheres traditionally used for Luminex assays and enables the simultaneous assessment of multiple surface EV markers with high sensitivity. This novel approach allows for the sensitive evaluation and in-depth analysis of specific EV populations in human plasma and cell culture supernatant (EV profiling) and offers several key improvements in the balance of sensitivity and specificity, a multiplexing capability and ease of analysis. Moreover, IEL is optimized for implementation with unprocessed plasma samples to bypass the requirement for EV enrichment and purification. This greatly simplifies the analysis of EV surface proteins in unmodified biofluids and renders it applicable for large population studies and biomarker discovery. Moreover, it eliminates an added bias due to potential selective EV loss introduced by diverse isolation techniques.

**Figure 1.**
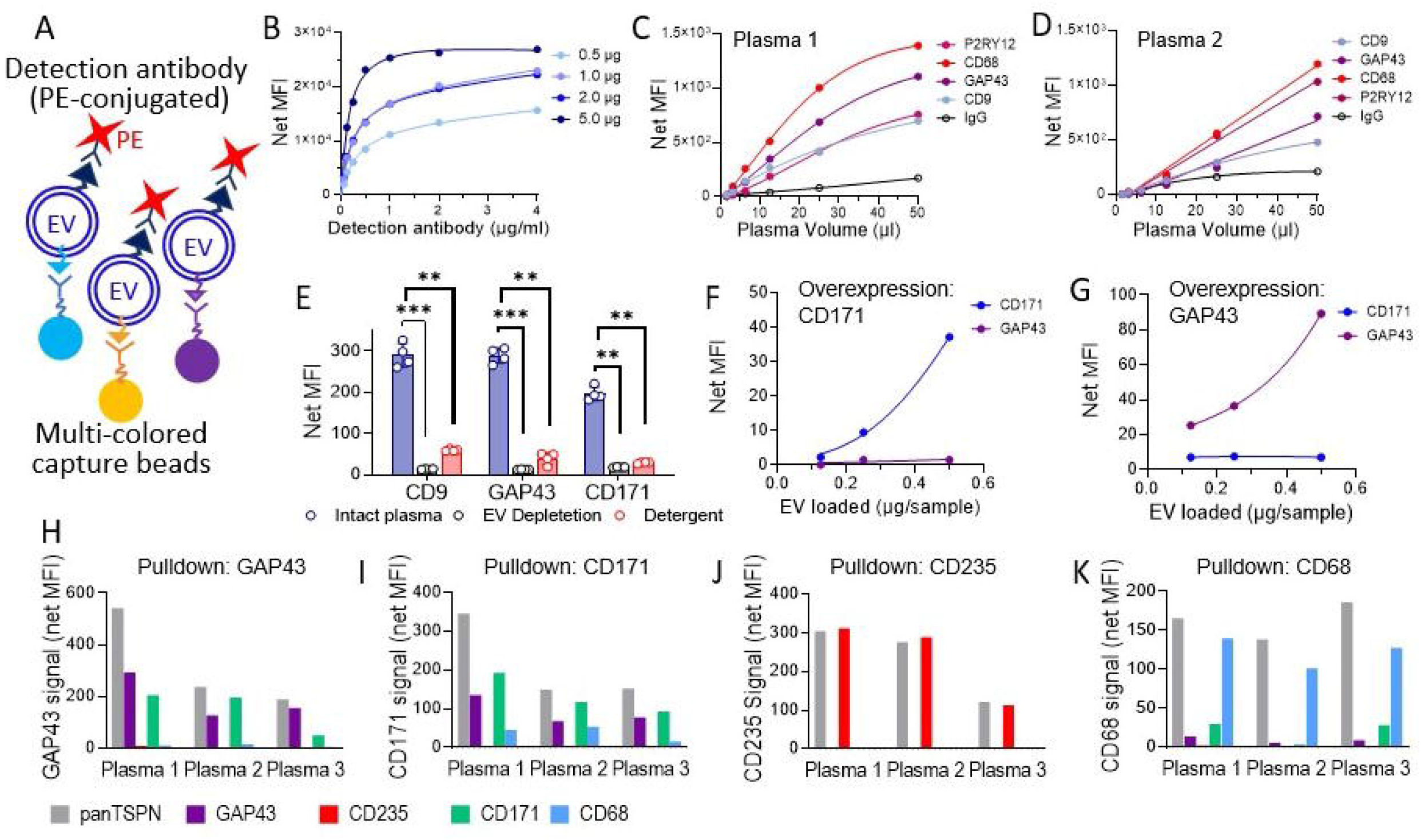
Optimization and testing of the beads for direct Luminex assessment of surface EV markers from blood plasma. **(A)** Schematic representation of a multiplexed bead-based sandwich assay (intact EV Luminex, IEL). Capture beads labeled with distinct fluorophores (MagPlex spheres, Luminex Corp.) are decorated with antibodies against specific surface antigens (color-coded in the diagram) and incubated with plasma or other EV-containing fluids. PE-tagged capture antibodies can vary, depending on the assay goals. Typically, a cocktail of EV markers, such as CD9, CD63 and CD81 is used. Note that capture and detection antigen should be co-presented on the surface of the same particle (connected by plasma membrane). **(B)** Antibody loading optimization for intact EV Luminex (IEL). MagPlex spheres were functionalized using increasing concentrations of mouse monoclonal antibodies against human CD9 (BioLegend, cat. no. 312102) in the indicated range of concentrations (0.5 – 5.0 μg/ml). Antibody loading was measured by incubation with increasing concentrations of PE-conjugated goat anti-mouse antibodies (BioLegend cat. No. 405307) and in a Luminex200 following standard procedure. **(C, D)** Quantitative detection of specific EV populations in two distinct plasma samples. MagPlex beads loaded with the indicated primary antibody (1 μg/ml) and 250 beads/sample were used to probe unprocessed human plasma (BioIVT). Plasma samples were diluted as indicated in 100 μl 1x PBS supplemented with 1%BSA (see Methods). Beads loaded with non-specific mouse IgG were used as a negative control. The detection was conducted using the cocktail of PE-conjugated panTSPN specific antibodies (see methods). Note a dose-dependent signal with clear linear range and sample-specific differences attributable to the unique representation of specific EV populations. **(E)** Total EV populations in the plasma were depleted prior to analysis, by polymer precipitation, or lysed using detergent-based buffer. All samples were probed with CD9 conjugated MagPlex spheres and the signal detected with panTSPN antibody cocktail. Note significantly lower signal following EV depletion or detergent lysis (**, P<0.001; ***, P<0.0001 by Tukey’s multiple comparison’s test). **(F, G)** EVs were isolated using ion exchange chromatography from cell culture media conditioned by HEK293 cells overexpressing neuron-specific antigens CD171 (F) and GAP43 (G). The EVs were diluted in 1x PBS supplemented with 1% BSA in 1 PBS and probed as indicated, with the beads conjugated with antibodies against human CD171 and GAP43, respectively. The detection was conducted with the panTSPN antibody cocktail. Note the lack of non-specific detection. **(H-K)** Plasma samples were analyzed using IEL beads for GAP43 (H), CD171 (I), CD235 (J) and CD68 (K), paired with the indicated detection antibodies (panTSPN cocktail, GAP43, CD171, CD68 and CD235, as shown).

Since the main principle of Luminex is similar to that of flow cytometry (Graham et al. 2019), we followed a recently published list of key instrumental and procedural controls recommended for the flow-based EV analysis (Welsh et al. 2020), including the demonstration of signal purge by EV depletion (through precipitation-based removal or detergent-based membrane lysis), a dilution-dependent signal reduction, and lack of signal with control capture antibody, in unstained control, or with negative control detection antibody. In the course of assay development, we were able to demonstrate the lack of signal from EVs of irrelevant cellular origin (i.e., no reactivity of neuronal EVs with antibody against an erythrocyte marker, and vice versa). Moreover, in agreement with previous findings, our method shows that plasma EVs carrying distinct cell type-specific markers also have different tetraspanin composition (e.g., CD9 is the most prevalent tetraspanin on the surface of erythrocytic EVs, but not on neuronal EVs). In addition, IEL analysis of plasma EVs fractionated by size exclusion chromatography showed that EVs positive for either neuronal markers (GAP43, CD171) or for the macrophage marker CD68 reside in fractions are distinct from EVs positive for the erythrocytic marker CD235a, suggesting differences in size distribution.

In conclusion, we have developed a novel, sensitive and robust methodology incorporating Luminex technology to define EV surface protein composition, supplementing existing EV characterization methods and enabling the rapid multifactorial analysis of tissue-specific EV biomarkers in health and disease.

## Material and Methods

### Preparation and characterization of IEL beads

Antibodies were conjugated to functionalized magnetic microspheres in desired luminescence range, functionalized with carboxyl groups (MagPlex^R^, Luminex Corp., Cat. No. MC1XXXX-01), using 1-Ethyl-3-(3-dimethylaminopropyl) carbodiimide (EDC) chemistry (He et al. 2007). Bead recovery/concentration after conjugation was determined in a Countess™ 3 FL Automated Cell counter (Thermo Fisher Scientific, Cat. No. A49866) using reusable chamber slides (Thermo Fisher Scientific, Cat. No. A25750). The beads were stored in a NeuroDex blocking buffer, which minimizes non-specific interactions at a minimum concentration of 1×10^6^/ml and for up to 3 months.

**To measure antibody loading**, IEL beads were resuspended at 5.0 × 10^6^/ml in Assay Diluent (NeuroDex) and loaded into 96-well black plates (BrandTech., Cat. No. 781671). A minimum of 200 beads/well and an equal volume of appropriate serially diluted PE-conjugated secondary antibodies was added in duplicate. Assay Diluent only was used as a background control. The staining was carried out for 60 min at room temperature in a Genie® Microplate Mixer/Shaker (600 RPM). Beads were washed twice in Wash Buffer 1 (NeuroDex) and twice more in Wash Buffer 2 (NeuroDex) resuspended in xMAP Sheath Fluid Plus (100 μl/well, Thermo Fisher Scientific, Cat. No. 4050021). The plates were analyzed in a Luminex200 plate reader, minimum 100 beads per condition.

### Preparation of biotinylated detection antibodies

Antibodies were biotinylated by overnight incubation at -4°C with EZ-link™ (Thermo Fisher Scientific,), at 20-fold molar excess, per manufacturer’s instructions. Excess reagent was removed using Zeba™ Spin Desalting Columns, 7K MWCO (Thermo Fisher Scientific, Cat. No. 89882).

### IEL procedure

50 μl of test sample diluted to desired concentration with Assay Diluent were placed in duplicates in the wells of a 96-well black plate. Assay Diluent only was used as a background control. Working IEL bead/microsphere suspensions were generated to yield 2.5-5.0 x 10^3^ beads/ml for each antibody/bead combination, for up to 7 IEL analytes, and 50 μl of working suspension was added to each well. The pull-down was carried out overnight (16-18 hours) at 4°C on a Genie microplate shaker (600 RPM). The plates were washed twice in Wash Buffer 1 (NeuroDex) and twice more in Wash Buffer 2 (NeuroDex) in a magnetic plate holder, and the beads were resuspended in Assay Diluent containing biotinylated detection antibody (1-2 μg/ml, 50 μl/well). After 2 hours incubation at room temperature (Genie shaker), the beads were washed 3 times in Wash Buffer 2 (NeuroDex), resuspended in 50 μl PBS containing Streptavidin-PE reagent (SAPE, 6 μg/ml) and incubated 20 min at room temperature. Following SAPE incubation, the beads were washed 3 times in Wash Buffer, resuspended in xMAP Sheath Fluid, and the plate was read on a Luminex200 reader.

### Size exclusion chromatography

EVs from the cell culture conditioned media were concentrated using a 100kDa MWCO Amicon concentrator (EMD Millipore, Cat. No. UFC910096), and plasma EVs were concentrated by precipitation with an optimized PEG reagent (NeuroDex). The concentrated samples (0.5 and 2 ml, as appropriate) were loaded onto SEC columns (IZON qEV Original, 35 nm and IZON qEV2, 35 nm, for conditioned media and plasma, respectively). Void volume and up to 25 fractions (0.5 and 2 ml, respectively) were collected, and protein concentration (A280 absorption) was measured in each fraction using a NanoDrop2000 spectrophotometer (Thermo Fisher Scientific).

### EV isolation from conditioned media

Conditioned media from induced pluripotent stem cell (iPSC)-derived neurons (120ml) was purchased from BrainXell. iPSCs had been differentiated into cortical neurons according to a proprietary protocol from BrainXell that does not include animal serum and thus avoids the risk of xenogeneic EV contamination. Conditioned media was collected across 4 weeks, where 50% of the media were collected each week. HEK293 cells were initially grown with media that contained 10% FBS; when the cells reached 50-70% confluence, the cells were transferred into EV-free, serum-free basal media, and the cells were incubated for an additional 48 hours. The media were collected, cleared by two centrifugation rounds (10 min at 3,000xg), and EVs were collected by ion exchange chromatography on DEAE Sephadex A-50 (20 mL) as described previously (Kosanovic et al. 2017). The unbound material was washed with equilibration buffer (0.05 M Tris–HCl, pH 7.6), followed by step elution with 0.05 M Tris–HCl, pH 7.6 0.25M NaCl, for weakly bound proteins and with 0.05 M TRIS–HCl buffer, pH 7.6, containing 0.5 M NaCl for EVs. Finally, the EVs were concentrated by dialysis against NeuroDex binding buffer, in a 100 kDa MWCO spin filtration unit (Amicon).

### Transmission Electron Microscopy

EV suspensions were fixed in 1% paraformaldehyde overnight. 4 μl of the suspension was loaded onto glow discharged copper mesh Formvar coated carbon grids and allowed to adsorb for approximately 30 seconds. The grids were briefly washed in double-distilled water, followed by staining with 2.5% aqueous Uranyl Acetate solution, and allowed to dry fully before imaging.

Grids were imaged using a FEI Morgagni transmission electron microscope (FEI, Hillsboro, OR) operating at 80 kV and equipped with a Nanosprint5 CMOS camera (AMT, Woburn, MA).

### Detergent treatment of crude EV fractions

Crude EV fractions generated by PEG-based precipitation (plasma EVs) or by centrifugation/ultrafiltration through 100 kDa MWCO Amicon spin filter (Millipore UFC901008) were supplemented with Triton X-100 (20% solution added to a final 2%) and incubated for 1-2 hours at room temperature or for 30 min at 50°C.

### RNA isolation and amplification

For RNA isolation, SEC fractions 2-5 and 9-14 were pooled and concentrated using a 100kDa MWCO Amicon filtration unit. RNA was isolated using the miRNeasy serum/plasma kit (Qiagen, Cat. No 217184) following the manufacturer’s instructions with modifications (treatment with RNase-free DNase). cDNA was generated using SuperScript^™^ VILO^™^ cDNA Synthesis Kit (Thermo Fisher Scientific, Cat. No: 11754050) according to the manufacturer’s instructions. The mRNA TaqMan kit and primers (Thermo Fisher Scientific, Cat. No. 44-445-56 and 4331182) are described in Table 3.

### Lipidomic analysis

Lipidomic profiling was performed using Ultra Performance Liquid Chromatography-Tandem Mass Spectrometry (UPLC-MSMS). Lipid extracts were prepared from live microsomes using a modified Bligh and Dyer method (Bligh and Dyer 1959), then spiked with appropriate internal standards, and analyzed using a platform comprising Agilent 1260 Infinity HPLC integrated to Agilent 6490A QQQ mass spectrometer controlled by Masshunter V 7.0 (Agilent Technologies, Santa Clara, CA). Glycerophospholipids and sphingolipids were separated with normal-phase HPLC as described before (Chan et al. 2012). An Agilent Zorbax Rx-Sil column (2.1 x 100 mm, 1.8 µm) maintained at 25°C was used under the following conditions: mobile phase A (chloroform: methanol: ammonium hydroxide, 89.9:10:0.1, v/v) and mobile phase B (chloroform: methanol: water: ammonium hydroxide, 55:39:5.9:0.1, v/v); 95% A for 2 min, decreased linearly to 30% A over 18 min and further decreased to 25% A over 3 min, before returning to 95% over 2 min and held for 6 min. Separation of sterols and glycerolipids was carried out on a reverse phase Agilent Zorbax Eclipse XDB-C18 column (4.6 x 100 mm, 3.5um) using an isocratic mobile phase, chloroform, methanol, 0.1 M ammonium acetate (25:25:1) at a flow rate of 300 μl/min.

Quantification of lipid species was accomplished using multiple reaction monitoring (MRM) transitions (Hsu et al. 2004; Guan et al. 2007; Chan et al. 2012) under both positive and negative ionization modes in conjunction with referencing of appropriate internal standards: PA 14:0/14:0, PC 14:0/14:0, PE 14:0/14:0, PG 15:0/15:0, PI 17:0/20:4, PS 14:0/14:0, BMP 14:0/14:0, APG 14:0/14:0, LPC 17:0, LPE 14:0, LPI 13:0, Cer d18:1/17:0, SM d18:1/12:0, dhSM d18:0/12:0, GalCer d18:1/12:0, GluCer d18:1/12:0, LacCer d18:1/12:0, D7-cholesterol, CE 17:0, MG 17:0, 4ME 16:0 diether DG, D5-TG 16:0/18:0/16:0 (Avanti Polar Lipids, Alabaster, AL). Lipid levels for each sample were calculated by summing up the total number of moles of all lipid species measured by all three LC-MS methodologies, and then normalizing that total to mol %. The final data are presented as mean mol % ± S.E.M.

## Results

### Intact EV Luminex allows specific detection of EV surface proteins in plasma samples

We generated IEL beads for the following surface antigens: CD9, a canonical EV marker; CD68, a macrophage marker; purinergic receptor P2RY12, a microglia marker; and neuronal marker axonal protein GAP43 (for specific capture antibodies, see Table 1). All antibodies were test-loaded at concentrations of 0.5, 1.0, 2.0 and 5.0 μg/ml, with bead concentration not exceeding 5×10^6^/ml, and loading was assessed by staining with serially diluted PE-conjugated secondary antibodies (Table 2) and measured in a Luminex200 reader (Thermo Fisher Scientific). Figure 1B shows CD9 loading as an example. For further IEL bead production, we chose antibody concentrations at the high end of a linear range, within which mean fluorescence intensity (MFI) increased proportional to the antibody concentration prior to saturation (1 μg/ml for the CD9 antibody and 1 - 5 μg/ml for other antibodies).

**Table 1:** Capture and detection antibody for intact EV Luminex.

| Antigen | Vendor | Cat. No. | Host/Type | Species specificity | Marker specificity |
| --- | --- | --- | --- | --- | --- |
| CD9 | BioLegend | 312102,<br>312112 | Mouse Monoclonal | human, simian | Pan-EV |
| CD63 | BioLegend | 353039,<br>353018 | Mouse Monoclonal | human, simian | Pan-EV |
| CD81 | BioLegend | 349502,<br>349514 | Mouse Monoclonal | human, simian | Pan-EV |
| CD171 | Thermo Fisher Scientific | 14-1719-82 | Mouse Monoclonal | human | Neuronal, gut |
| CD235a | BioLegend | 349502 | Mouse Monoclonal | human | Erythroid cells |
| GAP43 | Thermo Fisher Scientific | MA5-32256 | Rabbit monoclonal | Human, mouse, rat | Neuronal |
| P2RY12 | BioLegend | 848002 | Rat Monoclonal | Mouse, human | Microglia |

**Table 2:** secondary antibody to control IEL bead loading.

| Antibody | Vendor | Cat. No. |
| --- | --- | --- |
| PE Goat anti-mouse IgG | BioLegend | 312102 |
| PE Donkey anti-rabbit IgG | BioLegend | 406421 |
| PE Goat anti-rat IgG | BioLegend | 405406 |

IEL beads generated as in Figure 1A were tested for EV detection in unprocessed, serially diluted plasma samples from healthy donors (BioIVT), using a cocktail of biotinylated detection antibodies (CD9, CD63, and CD81) for tetraspanin detection (panTSPN, see Table 3 for detection antibodies). The analysis of two random unprocessed plasma samples using IEL beads conjugated with antibodies against specific targets showed strong dose-dependent signals with a linear assay range between 10 and 50 μl plasma volume (1/5 – 1/1 plasma dilutions, Fig. 1C, D. In contrast, IEL beads conjugated with non-specific IgG (a negative control) generated much lower signal, likely representative of non-specific binding (Fig. 1B, C). To ascertain that the signal can be attributed specifically to plasma EVs, we performed EV depletion prior to IEL using a NeuroDex-optimized PEG-based precipitation or detergent lysis (Triton-X100, 5% final concentration). In both cases, the signal was reduced by at least 80% (Fig. 1E). Finally, to demonstrate the specific capture of intended antigen, we used culture media conditioned by HEK293 cells overexpressing neuronal antigens GAP43 or CD171 (L1CAM; see Methods). Figure 1F shows that CD171-positive EVs in the culture media were recognized by the IEL beads appended with CD171 but not with GAP43 antibodies. Conversely, EVs from the media conditioned by GAP43-overexpressing cells were detected with GAP43 beads and not with CD171 IEL beads (Fig. 1G). Together, our results show that IEL detects a signal that is both EV-specific and antigen-specific in small volumes of unprocessed human plasma or conditioned media.

**Table 3:** Primers used in the study.

| Gene transcript | Gene ID | Vendor | Cat. No. |
| --- | --- | --- | --- |
| Neurogranin (NRGN) | Hs00382922_m1 | Thermo Fisher Scientific | 4331182 |
| SRY-Box Transcription Factor 1 (SOX1) | Hs01057642_s1 | Thermo Fisher Scientific | 4331182 |
| Oligodendrocyte Transcription Factor 1 (OLIG1) | Hs00744293_s1 | Thermo Fisher Scientific | 4331182 |
| Oligodendrocyte Transcription Factor 2 (OLIG2) | Hs00300164_s1 | Thermo Fisher Scientific | 4331182 |
| Hypocretin/Orexin (HCRT) | Hs01891339_s1 | Thermo Fisher Scientific | 4331182 |
| Glyceraldehyde-3-Phosphate Dehydrogenase (GAPDH) | Hs02786624_g1 | Thermo Fisher Scientific | 4331182 |
| Platelet Factor 4 (PF4) | Hs00427220_g1 | Thermo Fisher Scientific | 4331182 |
| Heterozygous Beta-Globin/ Hemoglobin (HBB) | Hs00758889_s1 | Thermo Fisher Scientific | 4331182 |

### IEL pulldown results in minimal cross-population contamination

Using IEL targeted towards specific EV populations, such as erythrocyte-derived or neuron-derived, we were able to demonstrate minimal cross-reactivity with antigens specific for unrelated cell types. Specifically, IEL beads decorated with antibodies against neuronal marker GAP43 were detected with high sensitivity using the panTSPN cocktail, GAP43 antibody and antibody against CD171 (Fig. 1H but not with the antibody against macrophage (CD68) and erythrocyte (CD235) markers. Similarly, CD171 pulldown resulted in IEL population detectable by panTSPN cocktail and by GAP43 antibody. Low positivity for CD68 is consistent with low-level CD171 expression by the circulating monocyte population (Fig. 1I. Conversely the EV population that was pulled down by CD235 IEL beads was detectable only with panTSPN and CD235 antibodies and lacked positivity for GAP43, CD171 and CD68 (Fig. 1J). Finally, CD68-positive EVs were detectable only with panTSPN, CD68, and to a lesser degree, CD171 antibody (Fig. 1K).

### Intact Exosome Luminex is suitable for the multiplexed assessment of EV surface proteins

One of the strongest advantages of the Luminex platform is its multiplexing capacity. To demonstrate the applicability of IEL for multiplexed measurements, we compared the sensitivity of multiplexed and single-bead IEL assays for multiple tissue-specific and canonical pan-EV surface markers including the erythrocyte marker CD235 (Glycophorin A), the macrophage marker CD68, EV markers CD63 and CD81 and neuronal markers GAP43 and CD171. Measurements in serial plasma dilutions showed a dose-dependent signal reduction for each analyte, with a clear linear range between 10-50 μl plasma input (Fig. 2A). To assess the potential reduction in sensitivity due to multiplexing, we generated dose/response curves for both single-bead and multiplexed assays for each analyte. Both types of assays generated similar curves within their linear range (Figure 2B-F). Moreover, measurements of CD9 in 25 random plasma samples produced significant differences between individual donors, but a strong correlation between single-bead and multiplexed assays (Figure 2G).

**Figure 2.**
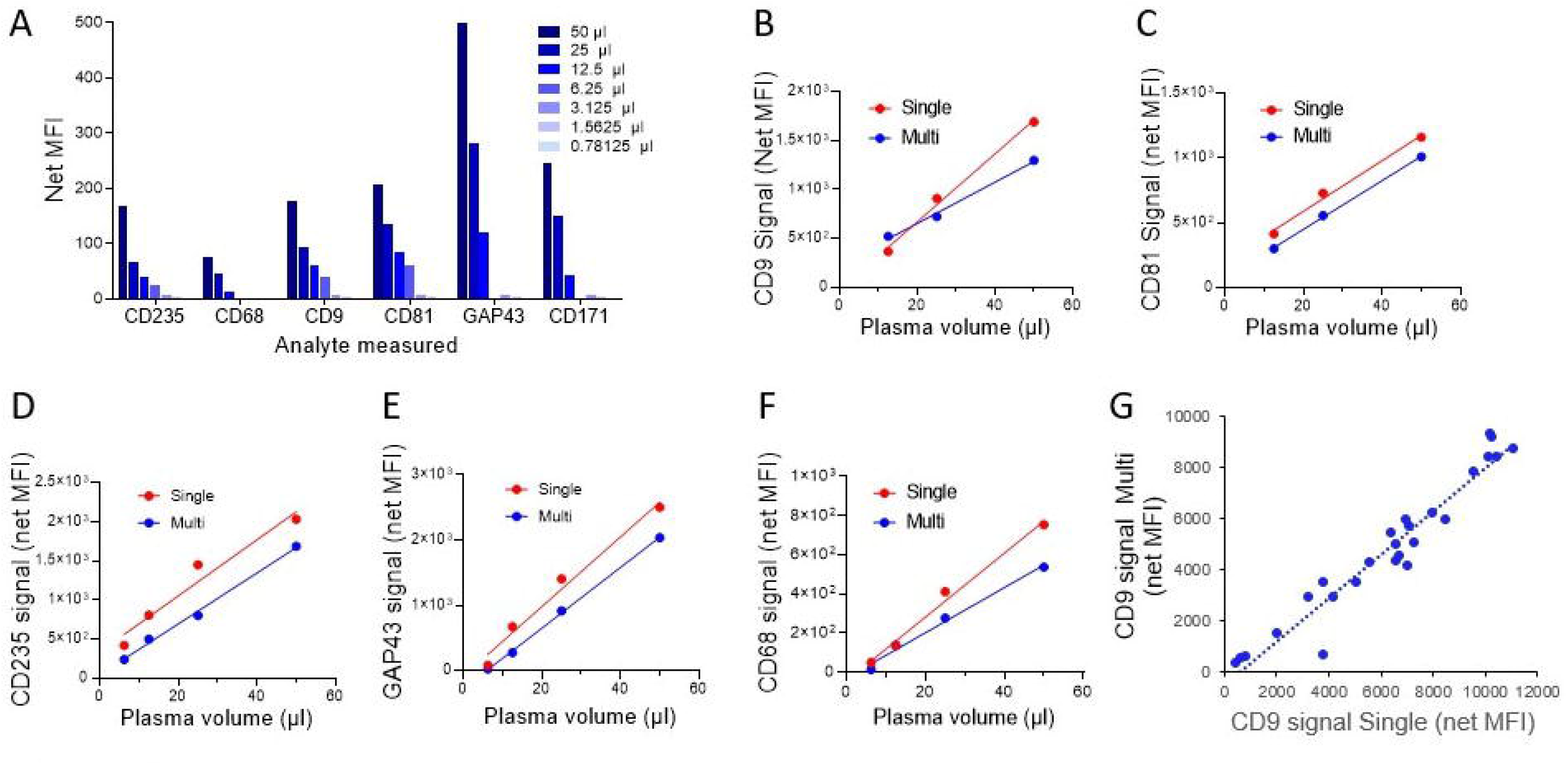
IEL Multiplexing capabilities enable simultaneous measurement of at least six surface EV proteins. **(A)** A multiplex assay for simultaneous detection of EV populations expressing major tetraspanins, CD9, CD63 and CD81, together with two neuron-specific markers (CD171 and GAP43) was used to probe the indicated dilutions of unprocessed human plasma. Note a clearly detectable signal in a linear range using 1-50 μl of blood plasma. **(B-G)** Unprocessed plasma samples were subjected to IEL with indicated antibodies in a multi- and single-plexed format and the values for each antibody plotted together for pairwise comparison. Note close absolute values and linearity in a similar dilution range (10-50 μl of plasma per measurement). **(H)** Twenty-five postmortem plasma samples were subjected to IEL in a multi- (6-pex) and single-plexed format (25 μl/sample) and the values for CD9 compared. Note strong correlation between individual values regardless between multiplexed and simplex measurements.

### Intact Exosome Luminex informs co-localization of EV surface molecules

To analyze potential differences in the expression of distinct surface protein by EVs of various cellular origins, we used multiple combination of IEL capture beads with detection antibodies. When GAP43 (neuron-specific) IEL beads were used in combination with either panTSPN or GAP43 antibody for detection (Figure 3A), we recovered similar signal strength, indicating that most GAP43 is EV-associated. Interestingly, when GAP43 IEL beads were applied with either CD9, or CD63, or CD81 capture antibodies, the signal generated with CD9 antibodies was significantly weaker (Figure 3B), suggesting that CD63 and CD81 are the two major tetraspanins expressed by GAP43-positive neuronal EVs. Interestingly, lower CD9 signal was also generated by neuronal cell culture-derived EVs compared to the signal for CD63 and CD81 (Figure 1H-K), but not by HEK293 cells (data not shown). When using CD235 IEL beads to capture erythrocytic EVs, only CD9, but not CD63 or CD81 detection antibodies, generated strong dose-dependent signal (Figure 3C), suggesting that the majority of CD235-positive EVs contain low levels of CD63 and CD81, but high levels of CD9. These differences in tetraspanin expression between EVs from different cell types suggest that differences in IEL signals are not solely attributable to differences in antibody affinity, but to differential surface presentation of specific antigens by different EV types. Our observation is consistent with previous studies (reviewed in (Thery et al. 2018; Witwer and Thery 2019)) which find CD9 expressed at high levels on erythrocyte derived EVs. Furthermore, EVs captured with CD9 IEL beads were positive for CD235 but to a lesser extent with GAP43 (Figure 3D), and EVs positive for CD63 or CD81 contained low level of CD235 signal compared to the signal detected with GAP43 antibody (Figure 3E, F). Together, these results demonstrate that IEL is a useful method to characterize EV surface protein composition, assess the colocalization of markers on the EV surface and potentially identify the divergent EV subpopulations.

**Figure 3.**
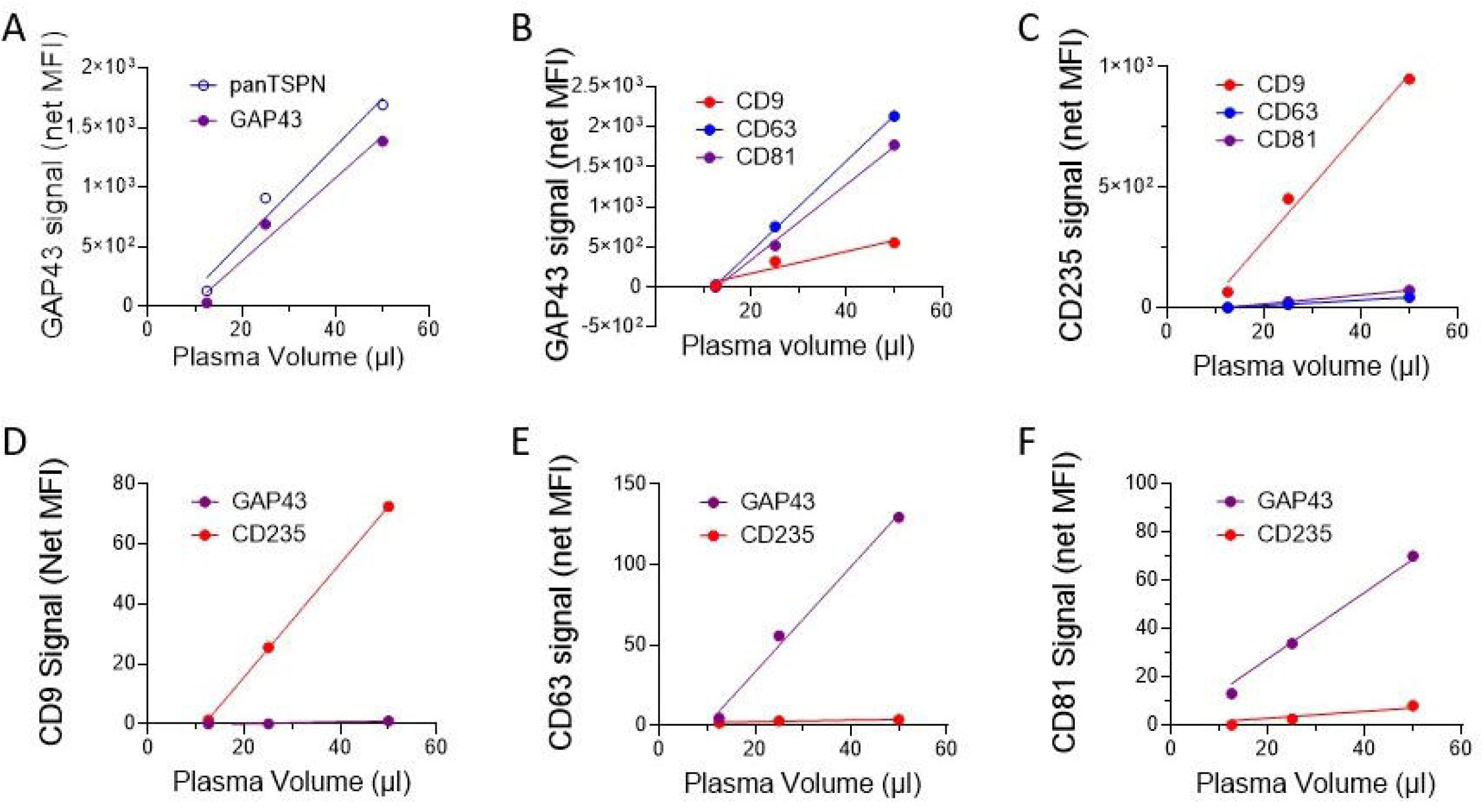
Intact EV Luminex reveals cell type specific differences between the tetraspanin profiles of distinct EVs populations in blood plasma. **(A)** Whole plasma (10-50 μl/sample) was subjected to IEL with antibodies against axonal marker, GAP43 followed by detection with PE-conjugated GAP43 antibodies or using panTSPN antibody cocktail. Similar values obtained for each detection method suggest that GAP43 in blood plasma is predominantly EV-associated. **(B)** Plasma IEL with GAP43 antibodies was followed by detection with individual TSPNs, CD9, CD63 and CD81, respectively. Note lower levels of CD9 in GAP43-positive EVs. **(C)** IEL for erythrocyte EVs using a known marker CD235 glycophorin was followed by detection with individual TSPNs, CD9, CD63 and CD81 antibodies, respectively. Note the high level of CD9 expression in CD235-positive EVs and negligible levels of CD63 and CD81. **(D)** IEL for one of the EV markers, CD9 was followed by detection with CD235 and GAP43 antibodies. Note low levels of GAP43 and strong positivity for CD235. **(E, F)** IEL for CD63 (E), CD81 (F) followed by GAP43 and CD235 detection shows strong positivity for GAP43 and the lack of detectable association with CD235.

### Intact EV Luminex combined with size exclusion chromatography uncovers cell-specific variability in fraction recovery

To further characterize the nature of the signal registered by IEL, we performed size exclusion chromatography (SEC) of plasma EVs followed by IEL analysis of collected fractions. To ensure that signal is generated by the EVs, we first used only the panTSPN detection cocktail. Plasma EVs were concentrated by PEG precipitation (see methods), resuspended in PBS, and loaded onto pre-calibrated qEV2 columns (IZON Sciences). Several void fractions (negative numbers) and all eluted fractions were analyzed by IEL with beads conjugated to antibodies against CD9, CD235, CD68, GAP43 and CD171. The combination of CD9 IEL beads with the panTSPN detection cocktail produced the expected signal that peaked between fractions 2-5 and trailed to fraction 9 (Fig. 4A). CD235-positive EVs detected with the panTSPN cocktail also peaked at fraction 3, but with a narrow peak suggestive of a tighter size range (Figure 4B). Unexpectedly, IEL beads for CD68, a marker of tissue macrophages, and for the two neuronal markers, GAP43 and CD171, while detectable with the same panTSPN cocktail, yielded strong signals in fractions 11-18, 8-15 and 11-18, respectively (Figure 4C-E), suggesting smaller particle sizes. Indeed, TEM analysis of pooled early and late fractions (qEV Original, IZON Sciences) showed spherical particles of 75-125 nm. in diameter in fractions 2-5 (Fig. 4F, left) and smaller particles in fractions 8-14 ranging between 15 and 25 nm in diameter (Fig. 4F, right).

**Figure 4.**
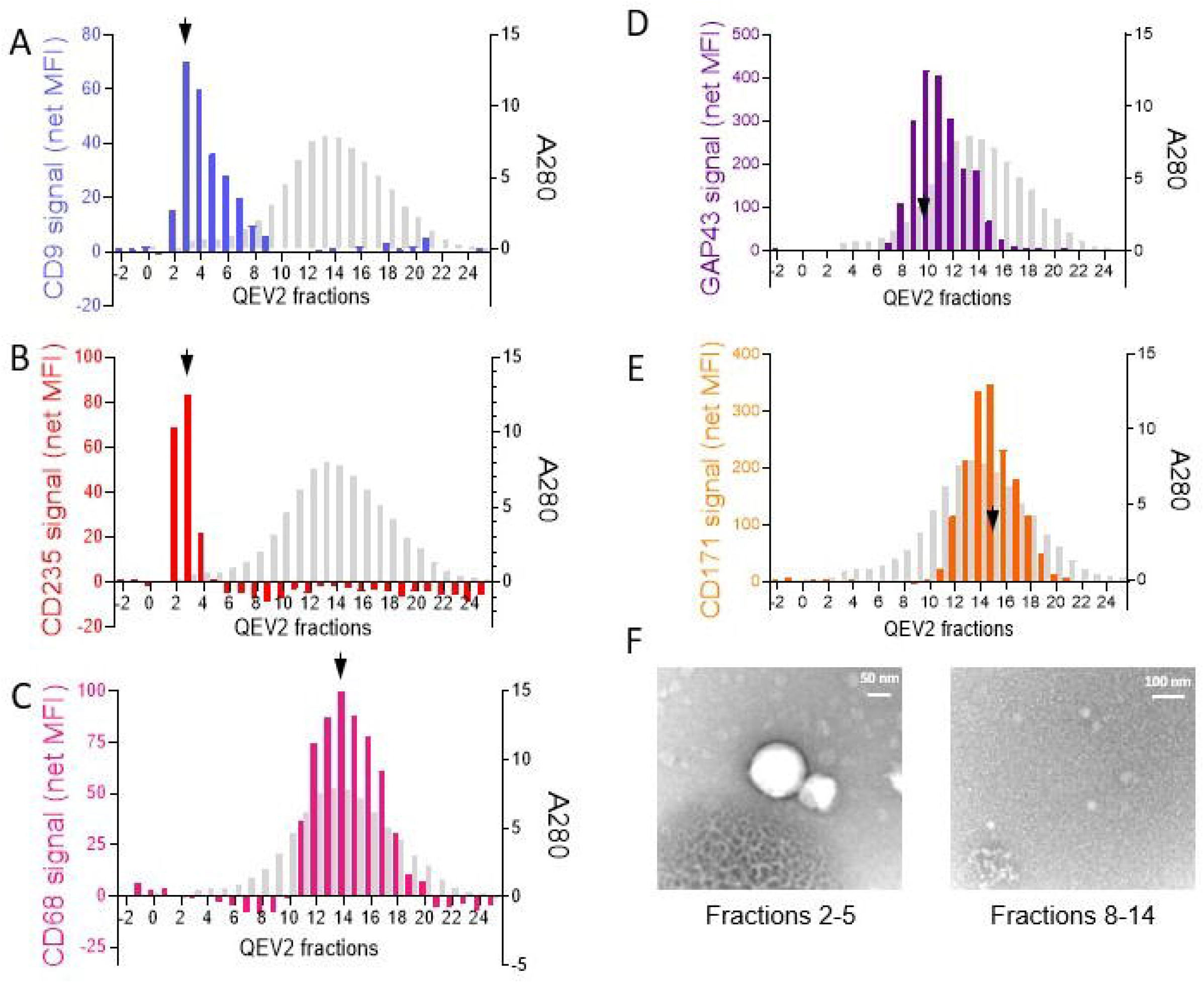
Intact EV Luminex reveals size differences between plasma EV populations positive for distinct cell type specific antigens. EVs from 10 ml plasma were precipitated using PEG-based approach, resuspended in 2 ml PBS, loaded onto QEV2 35nm column (IZON Sciences) and 2 ml fractions were collected following the manufacturer’s instructions. Protein concentration (A280) was measured in each fraction, including 3 void fractions (denoted -2, -1 and 0) and 50 μl aliquots analyzed by IEL as shown. The fractions with the peak concentrations of specific EVs (detected with panTSPN antibody cocktail) are indicated by arrows. **(A)** IEL for CD9, detection with panTSPN antibody cocktail. Protein elution is shown in grey. **(B)** IEL for erythrocyte marker CD235, detection with panTSPN antibody cocktail. **(C)** IEL for macrophage marker, CD68, detection with panTSPN antibody cocktail. **(D)** IEL for axonal marker, GAP43, detection with panTSPN antibody cocktail. **(E)** IEL for neuronal marker, CD171, detection with panTSPN antibody cocktail. Note that CD9-positive and CD235 positive EVs collect in early fractions, consistent with the average diameter of 150-200 nm, while the peaks of EVs positive for CD68, CD171 and GAP43 are consistent with diameters of 30-50 nm. and below. **(F)** Pooled, concentrated fractions 2-5 and 8-14 (qEV Original, IZON Sciences) were loaded onto microgrids, stained with aqueous uranyl acetate, and analyzed by transmission electron microscopy.

To further characterize the origin of these late fraction signals, we repeated the analysis, using GAP43 IEL beads followed by detection with panTSPN antibody cocktail or with specific TSPN antibodies (Figure 5A-D). Interestingly, while panTSPN antibody yielded strong positive signal in fractions 7-15, (Figure 5A), the detection of CD9 alone resulted in a weak (<50 net MFI) positive signal in fractions 11-16. CD63 and CD81 detection both yielded much stronger signals in fractions 9-15 and 7-16, respectively (Fig. 5C, D). In agreement, CD9 IEL beads in combination with GAP43 detection antibody produced a weak signal in fractions 10-16, while CD63 and CD81 IEL beads combined with GAP43 detection antibody yielded signal in fractions above 10.

**Figure 5.**
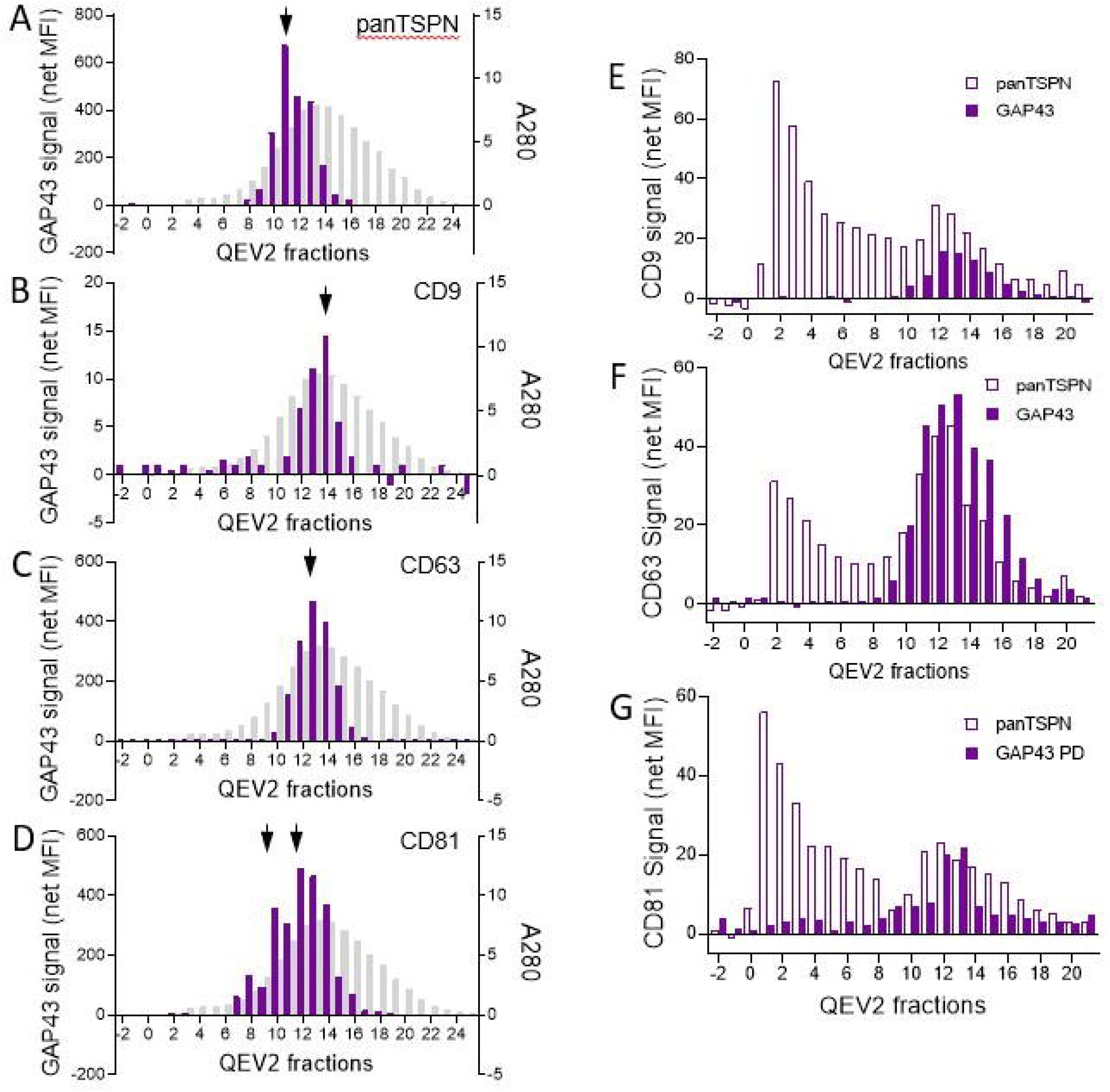
Intact EV Luminex reveals a continuum of sizes amongst tetraspanin-positive structures (EVs) in blood plasma. **(A-D)** Fractions collected from QEV2 column were subjected to IEL for axonal marker GAP43 and probed with the pan-TSPN antibody cocktail (A) or with the antibodies against CD8 (B), CD63 (C) and CD81 (D), respectively. Note that peak concentrations are observed in different fractions for distinct tetraspanins, covering a broad range of sizes. **(E-G)** QEV2 fractions were subjected to IEL for CD9 (E), CD63 (F) and CD81(G), and probed with antibodies against axonal marker, GAP43, or with panTSPN antibody cocktail, as shown. Note a broad range of sizes for all analytes and detection antibody combinations.

To ascertain whether the late-fraction signals detectable with pan-TSPN antibodies can be attributed to EVs, we performed fractionation of the crude EV preparation that was subjected to pre-treatment with detergent to disrupt the association, via plasma membrane, between the EV and cell type-specific markers in accordance with MISEV guidelines (Thery et al. 2018; Welsh et al. 2020). Treatment with Triton-X100 at 50°C resulted in significant reduction of the signal in early fractions positive for CD9, CD235 and CD63 (Fig. 6A-C). The same treatment significantly reduced the GAP43 signal in late fractions, and caused the shift of CD63 and CD235 signals towards late fractions (Fig. 6C, D). The latter may reflect the closure of residual EV membranes following detergent treatment causing the formation of smaller vesicles. Another potential reason for the reduced detergent sensitivity of smaller EVs is their distinct lipid composition, such as increased lipid raft content in the EV membranes (Sonnino and Prinetti 2013). This hypothesis is supported by lipid content analysis (shown below).

**Figure 6:**
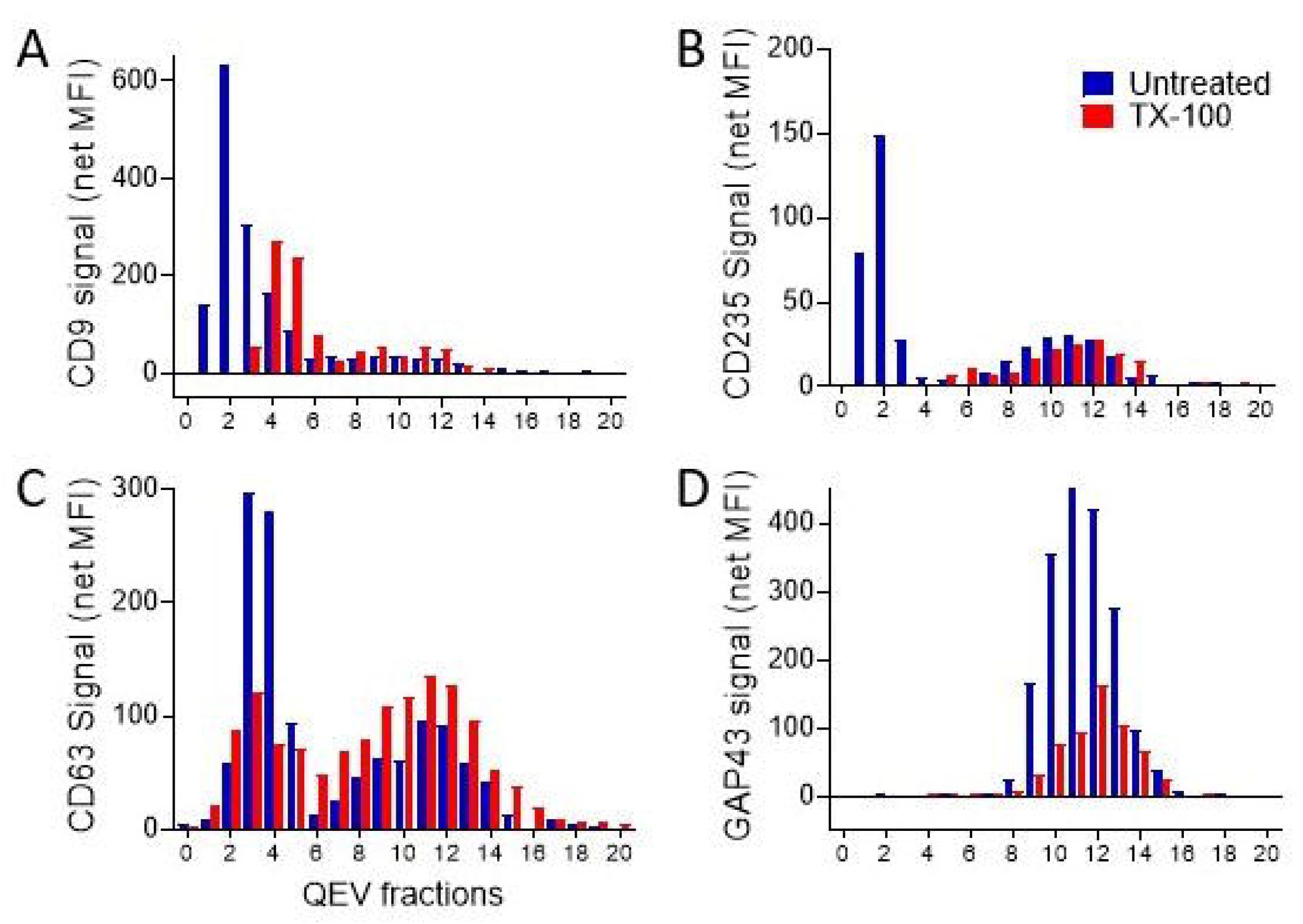
Plasma membrane disruption by Triton X-100 pre-treatment of plasma samples diminishes IEL signal. Plasma samples were concentrated by PEG-based precipitation, reconstituted in 1x PBS. The samples were then pre-treated with TX-100 (red bars) or sham buffer (blue bars) and subjected to size-exclusion chromatography (IZON qEV2 column). The fractions were then subjected to IEL with **(A)** CD9, **(B)** CD235, **(C)** CD63 and **(D)** GAP43 beads paired with panTSPN detection cocktail. Note diminished signal upon of TX-100 pre-treatment, for both early and late fractions.

Interestingly, IEL beads for CD63 and CD81, in addition to strong peaks in early fractions, generated dispersed positive signals in later fractions (Figure 5E-G). These results are consistent with previous findings, where CD63 and CD81 detection in qEV fractions generated a continuous signal in early and late SEC fractions (Ter-Ovanesyan et al. 2021). While this finding was interpreted as contamination, and similar distribution was not observed in cell culture media (Fig. 7), it may also reflect a broader size distribution of true EVs in blood plasma.

**Figure 7:**
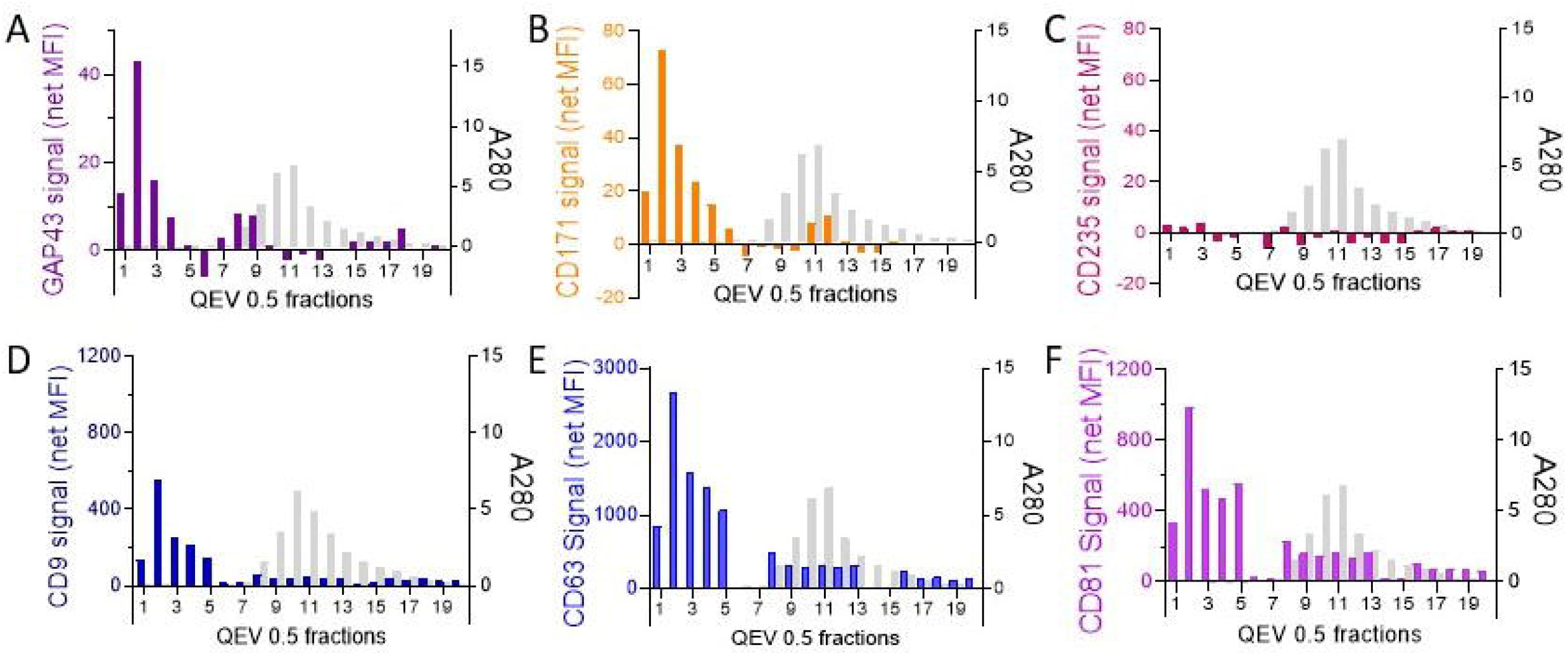
IEL analysis of neuronal cell conditioned media. Serum-free media conditioned by iPSC-derived neurons (BrainXell, 10 ml) was collected, cleared by filtration via Amicon spin filter (100 kDa MWCO) reconstituted in 0.5 ml PBS, and processed by SEC using QEV original column (35 nm, ZON Sciences). Fractions (0.5 ml) were collected according to the manufacturer’s instructions and protein concentration (A280) determined in each fraction (gray bars). All fractions (1-20, 50 μl per sample) were subjected to IEL with beads for **(A)** GAP43, **(B)** CD171 and **(C)** CD235 followed by detection with panTSPN antibody cocktail. Note the lack of CD235-positive EVs in the media conditioned by iPSC neurons (specificity control). **(D-F)** IEL of the QEV fractions with beads for EV markers, CD9 (D), CD63 (E) and CD81 (F). Note for each marker, except CD235, a secondary smaller peak or disperse signal in the late fractions.

### SEC late fractions contain mRNA and lipids typical of EVs

Previously it has been postulated that the late fractions generated by size exclusion do not contain EVs; however, TSPN-positive signal detected by IEL in the later fractions suggests the possibility of EV presence. To further confirm the EV nature of the signal, we examined RNA and lipid content of the pooled fractions 2-5 and 8-15. EVs are the main carrier of cell-free mRNA (Jose 2015; O’Brien et al. 2020). Early fractions (1-6) and late fractions (8-15) were combined and concentrated by spin-filtration (using 100 kDa MWCO Amicon units), prior to RNA extraction. Multiple cell-specific mRNAs were measured by qPCR with TaqMan primers (Table 3) using an amplicon-based approach, (the results are summarized in Table 4A-C). An approximately 2-fold lower GAPDH content in fractions 8-15 indicates higher overall RNA levels in fractions 1-6, consistent with the predicted larger EV size; however, several mRNA were clearly detectable in fractions 8-15. Moreover, normalization to the housekeeping transcript (GAPDH), revealed 2 to 4-fold enrichment in neuron-specific transcripts, such as neurogranin (NRGN) and Orexin (HTCR) as well as oligodendrocyte markers SOX1 and OLIG2 (Table 4A). This is consistent with high content of neuron-specific protein markers, e.g., GAP43 and CD171 (see above). In addition, the ratios between neuron-specific mRNA (NRGN) to erythrocytic and platelet-specific mRNA (HBB and PF4, respectively) are significantly higher in the later fractions (Tables 4A, B). Together, these findings suggest distinct cellular origins of the EV populations in the early and late fractions, and that in plasma, the majority of EVs of neuronal origin are detected in the lower size range.

**Table 4A:** mRNA enrichment in pooled SEC fractions (normalized to GAPDH)

| Gene transcript | $\Delta C_t$ | | Late/early ratio |
| --- | --- | --- | --- |
|  | Early fractions | Late fractions |  |
| NRGN | 2.362234 | 0.407699 | 3.875911 |
| SOX1 | 2.589385 | 0.71876812 | 3.656889 |
| OLIG1 | 2.192837 | 1.758674622 | 1.351127 |
| OLIG2 | -2.35433 | -3.76834 | 2.664775 |

**Table 4B:** Neuronal and erythrocytic mRNA content in pooled SEC fractions.

| Gene transcript | Ct | | ΔCT (to HBB) | | Ratio to HBB transcript | |
| --- | --- | --- | --- | --- | --- | --- |
|  | Early fractions | Late fractions | Early fractions | Late fractions | Early fractions | Late fractions |
| NRGN | 22.99162312 | 23.04724693 | -5.61276 | -8.06409931 | 48.93386414 | 267.6306081 |
| SOX1 | 28.92022133 | 28.06206989 | 0.315836906 | -3.04927635 | 0.803384813 | 8.277966166 |
| OLIG1 | 28.52367401 | 29.10197639 | -0.08071041 | -2.00936985 | 1.057538665 | 4.026063286 |
| OLIG2 | 23.97650909 | 23.57496071 | -4.62787532 | -7.53638553 | 24.7246009 | 185.6427979 |
| HBB | 28.60438442 | 31.11134624 | - | - |  |  |

**Table 4C:** Neuronal and platelet mRNA content in pooled SEC fractions.

| Gene transcript | Ct | | ΔCT (to HBB) | | Ratio to PF4 transcript | |
| --- | --- | --- | --- | --- | --- | --- |
|  | Early fractions | Late fractions | Early fractions | Late fractions | Early fractions | Late fractions |
| NRGN | 22.99162312 | 23.04724693 | -9.07498531 | -13.4121246 | 539.3153602 | 10900.63556 |
| SOX1 | 28.92022133 | 28.06206989 | -3.1463871 | -8.39730167 | 8.854354284 | 337.1628269 |
| OLIG1 | 28.52367401 | 29.10197639 | -3.54293441 | -7.35739517 | 11.65546306 | 163.9821729 |
| OLIG2 | 23.97650909 | 23.57496071 | -8.09009933 | -12.8844108 | 272.4975285 | 7561.259528 |
| PF4 | 32.06660843 | 36.45937157 | - | - | - | - |

To further confirm the EV nature of the signal detected by IEL in the late SEC fractions, pooled fractions 1-6 and 8-15 were subjected to lipidomic analysis by Ultra Performance Liquid Chromatography-Tandem Mass Spectrometry (UPLC-MSMS). The lipid composition of fractions 8-15 was consistent with the published data generated by lipidomic analysis of EV membranes (Yanez-Mo et al. 2015; Skotland et al. 2020), wherein for the major classes of lipids (cholesterol, phosphatidylcholine and PC ethers, phosphatidylserine, phosphatidyl-ethanolamine and PE ethers, diglycerides, phosphoglycerates, phosphatidic acid, phosphatidyl inositol, ceramide and lactoceramide), the measured percentage of total lipid content in the late and early EV fractions, was within the previously observed range. Interestingly, we observed significant differences between the lipid content of the early (1-6) and late (8-15) fractions (Table 5 and Fig. 8). First, free cholesterol content was approximately 2-fold lower in the late fractions, likely due to sequestration in protein complexes involved in cholesterol transport in circulation (Cockcroft 2021; Zanotti et al. 2021). This was consistent with 2 to 3-fold higher amounts of phosphatidylethanolamine (PE), PE esters, phosphatidylcholine, and phosphatidylserine in fractions 8-15 (Table 5). On the other hand, significant 2.5-fold increase in sphingomyelin (SM) content and increased SM/cholesterol ratio, point to increased proportion of lipid raft domains (Simons and Ehehalt 2002; Ouweneel et al. 2020; Shaw et al. 2021). Given relative resistance to Triton-X100, these observations suggest higher lipid raft content in smaller EVs found in the late SEC fractions, compared to canonical EVs eluted in early fractions (Sonnino and Prinetti 2013; Pera et al. 2017).

**Table 5:** Lipid content of the late and early pooled SEC fractions.

| Lipid species | % Total lipid |  |  |  | Late/<br>Early<br>ratio | P value<br>(T-test) |
| --- | --- | --- | --- | --- | --- | --- |
|  | Early fractions |  | Late fractions |  |  |  |
|  | Average | Std | Average | Std |  |  |
| Cholesterol | 58.84417 | 12.78809 | 31.17807 | 1.001808 | 0.524965 | 0.013858 |
| Phosphatidylethanolamine | 7.185072 | 1.791263 | 18.09836 | 2.149141 | 2.518883 | 0.006269 |
| Phosphatidylethanolamine esters | 6.556616 | 1.639689 | 16.15601 | 1.432092 | 2.464078 | 0.010448 |
| Phosphatidylcholine | 5.06497 | 3.500439 | 16.9832 | 1.808727 | 3.35308 | 0.001742 |
| Phosphatidylserine | 2.803951 | 0.032804 | 5.460818 | 0.094241 | 1.947544 | 3.71E-05 |
| Dihydrosphingomyelin | 2.661473 | 0.419849 | 0.561044 | 0.017827 | 0.210802 | 0.04453 |
| Sphingomyelin | 2.066918 | 0.116012 | 5.86615 | 0.229976 | 2.838114 | 0.000206 |
| Sulfatide | 1.044329 | 0.121476 | 0.423145 | 0.051607 | 0.405184 | 0.030678 |
| Ceramide | 1.015121 | 0.101341 | 0.590466 | 0.026987 | 0.58167 | 0.046919 |
| Phosphatidylinositol | 1.009969 | 0.116646 | 2.007596 | 0.204886 | 1.987779 | 0.003097 |
| Lactosyl-ceramide | 0.804356 | 0.202362 | 0.153533 | 0.017996 | 0.190877 | 0.068087 |
| Phospho-ceramide | 0.344956 | 0.030619 | 1.191594 | 0.100097 | 3.454337 | 0.001015 |
| Monohexosyl-ceramide | 0.33583 | 0.01688 | 0.285899 | 0.019925 | 0.851319 | 0.034268 |
| Lyso-phosphatidyl ethanolamine | 0.222608 | 0.028278 | 0.142429 | 0.020116 | 0.63982 | 0.046703 |
| Monoglycerides | 0.177055 | 0.004027 | 0.640425 | 0.240593 | 3.617098 | 0.039629 |
| Cholesterol esters | 0.17601 | 0.009814 | 0.263788 | 0.045196 | 1.498709 | 0.035008 |
| Lyso-phosphatidyl choline | 0.15395 | 0.015923 | 0.087663 | 0.007112 | 0.569427 | 0.038256 |
| Lyso-phosphatidylethanolamine | 0.145211 | 0.020561 | 0.068489 | 0.011437 | 0.471648 | 0.03779 |
| Diglycerides | 0.1258 | 0.045601 | 0.058413 | 0.009868 | 0.464329 | 0.138111 |

**Figure 8:**
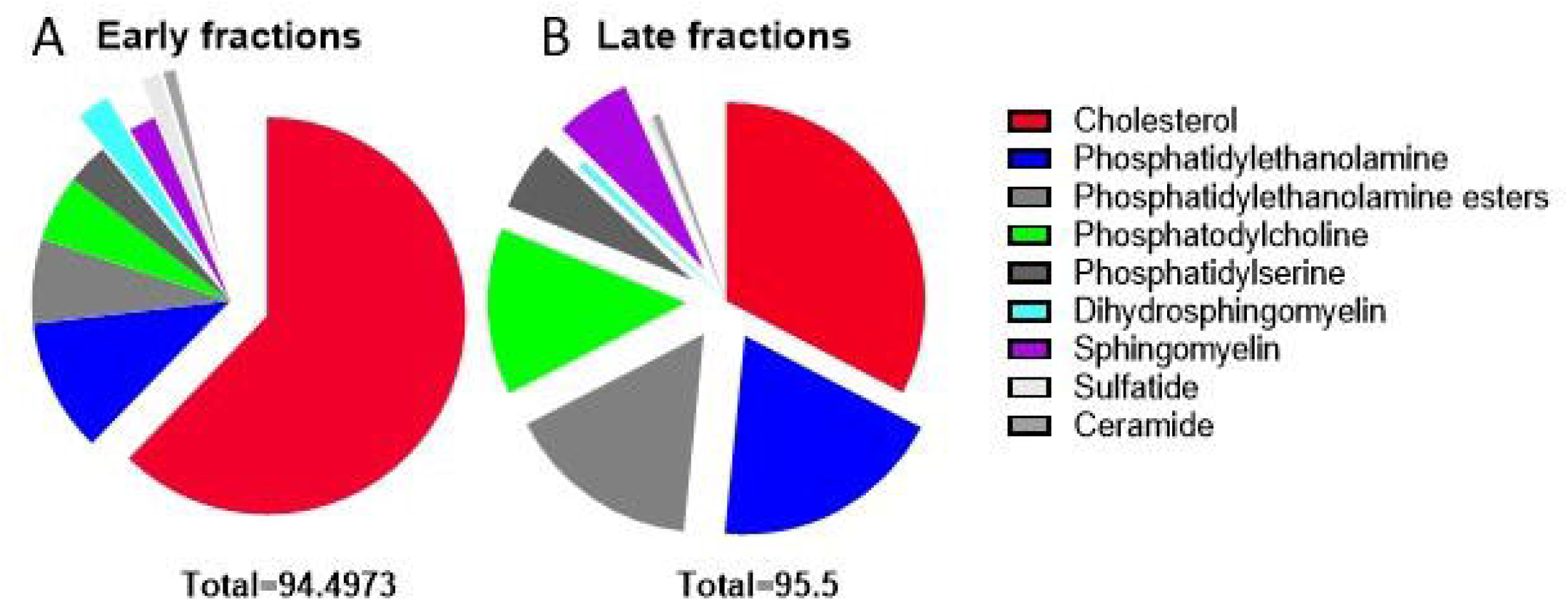
Lipid composition of late and early SEC fractions. Plasma EVs were concentrated as described in Methods and processed by SEC (qEV2, IZON, Sciences). Fractions 2-5 and 8-13 were pooled, concentrated using Amicon spin filter (100kDA MWCO) and subjected to lipidomic analysis (see Methods). Each fraction was analyzed in biological triplicate. The pie charts show proportional contribution (% of total lipid) for the indicated specific lipid classes in the early **(A)** and late fractions **(B)** is shown. “Exploding” sectors indicate lipids with higher proportional content. For minor lipid constituents (average content in early fractions < 0.1%), see Table 6.

Together, these observations strengthen the interpretation wherein the signal detected by IEL in the late SEC fractions represents diverse EV subpopulations rather than protein contamination.

### IEL analysis of SEC fractions of neuronal conditioned media detects EV peak in early fractions

To compare EV size distribution in blood plasma and in tissue culture media, EVs from conditioned media of iPS-derived cortical neurons (BrainXell) were analyzed. Conditioned media were concentrated by ultrafiltration (see methods), was subjected to SEC (qEV Original, IZON Sciences), the fractions were collected and analyzed alternatively by IEL with beads for GAP43, CD171 and CD235 combined with panTSPN detection (Figure 7A-C). Consistent with their neuronal origin, the EVs in early fractions showed strong positivity for GAP43 and CD171 (Figure 7A, B); however, approximately 10% of the total signal localized to the late fractions. In agreement with their neuronal differentiation, IEL analysis with CD235 beads yielded no detectable signal in any of the fractions (Fig. 8C), further confirming the detection specificity.

In addition, the same qEV fractions were analyzed using CD9, CD63 and CD81 IEL beads followed by GAP43 antibody detection (Figure 8D-F). All tetraspanins eluted with a similar pattern, a larger peak in early fractions and a lower dispersed signal in late fractions, suggesting a predominance of larger EVs in cell culture condition media. The CD63 and CD81 signals in the EVs from iPSC-derived neurons were by an order of magnitude higher than the signal for CD9, suggesting that while EV size distribution in neuronal cell culture is different than in blood plasma, their TSPN composition remains similar, with relative paucity of CD9-positive EVs.

## Discussion

Our goal was to create a reliable, easy to use assay for rapid multifactorial profiling of EVs in biological fluids. EVs are released from most tissues/organs and are found at relatively high concentrations in a variety of biofluids. Since EVs offer a molecular snapshot of their cell of origin, their promise as an emerging diagnostic and therapeutic platform is hard to overestimate (Simpson et al. 2009; He et al. 2018; Hunter and Dhaun 2020; Alberro et al. 2021; Heydari et al. 2021; Liu et al. 2021; Thakur et al. 2021; Trino et al. 2021). However, EVs isolation and characterization are hindered by several obstacles, the most daunting of which is extreme heterogeneity that extends to EV size, composition, and surface antigen profiles, which limits their implementation in clinical practice (Quiroz-Baez et al. 2020; Bordanaba-Florit et al. 2021; Claridge et al. 2021; Cloet et al. 2021; Singh et al. 2021). Characterization of EV surface proteins, an integral part of MISEV guidelines and requirements for EV studies, is a valuable approach, which may aid the identification of diverse subpopulations in complex EV pools found in biofluids. Here, we describe successful development of a straightforward and robust technique for simultaneous detection and assessment of EV surface proteins in a multiplexed assay.

This novel method, based on a simple sandwich principle, is applied to intact EVs, with two antibodies directed against distinct proteins presumably connected by the EV membrane. This concept has been repeatedly exploited for EVs analysis, in assays ranging from ELISA to microfluidic devices. Here we pair this approach with the Luminex platform, which advantageously combines high sensitivity with the ease of multiplexing. Moreover, the method is optimized for the use with unprocessed plasma, thus bypassing lengthy and cumbersome steps of EV isolation, enrichment, and purification.

Assay validation according to the published guidelines for EV flow cytometry analysis and FDA guidelines for immunoassay development requires the linearity of signal, proportional to the concentration of detection antibody. We observed linearity across 16-fold antibody and plasma dilutions, within 0.07-2ug/ml antibody concentration (R=0.9) and 8 to 33-fold plasma dilution, with the range variance dependent on plasma specimen and the analyte measured. While this observed linearity range is acceptable, it is relatively narrow for some antibody and plasma samples, suggesting the need for further optimization of working dilutions. The assay specificity requirements for the platform are twofold: selectivity for EVs and specificity of the antibodies against the chosen surface protein(s). Selectivity to EVs was demonstrated by EV depletion using PEG precipitation or by EV lysis with mild detergent. In both cases we observed significant (over 50%) reduction in the measured signal. Antibody specificity was assessed by comparison with non-specific, isotype-matched control antibodies (IgG) and by combining antibodies presumably specific for divergent EV subpopulations, such as erythrocytic, neuronal or macrophage-derived. Indeed, IEL beads appended with antibodies against CD235 (a specific erythrocyte marker) yielded clear signal upon detection with tetraspanin antibody cocktail (a combination of common EV markers), but not with antibodies against neuronal markers like CD171 and GAP43 or against a macrophage marker, CD68 and vice versa. These results further demonstrate the assay specificity for both EVs and for chosen analytes, in this case population-specific EV surface proteins. Moreover, it shows that our novel IEL method can be used to assess co-localization of EV surface proteins. This feature can be exploited in the future for analysis and characterization of EV subtypes in blood plasma as well as in cell culture media. The capacity for selective detection of either CD171 or GAP43, but not CD235 on EVs produced by HEK293 overexpressing either one or the other of these two neuronal proteins provides yet another indication of specificity. Similarly, EVs isolated from the media conditioned by iPSC-derived neurons yielded clearly detectable signals when probed with IEL beads conjugated with antibodies against neuronal (CD171, GAP43) and pan-EV (CD63 and CD81) antigens, but not with antibodies against erythrocytic marker CD235. The presence of CD171 on neuronal EVs generated in cell culture is in keeping with previous reports (Faure et al. 2006; Lachenal et al. 2011; Shi et al. 2014; Norman et al. 2021) and detection of GAP43/neuromodulin (Pfenninger et al. 1991), is another neuron-specific membrane protein, corroborates the assay specificity. Together, our results clearly support assay specificity and linearity.

Following IEL qualification as a single assay, we analyzed its utility for multiplexed analysis, a capacity that is particularly important to generate complex sets of data from the low volumes of plasma samples collected in clinical studies. Both linearity and detection range were similar between single and multiplexed assays for at least six EV markers (CD235, CD68, CD9, CD81, CD171 and GAP43), in an assay that combines up to 6 analytes. Moreover, the analysis of plasma samples from 20 healthy volunteers showed considerable variability of tetraspanin measurements between individuals, with strong (R^2^=0.93) correlation between the values generated using single and multiplexed assays. In the future, this assay may enable population-wide studies of tissue-specific EV species in blood plasma (EV profiling).

For additional verification of the EV nature of the signal(s) measured by the novel IEL assay, it was coupled with a standardized EV isolation procedure using size exclusion chromatography with pre-calibrated columns manufactured by IZON Sciences (Monguio-Tortajada et al. 2019; Kurian et al. 2021; Stam et al. 2021). Interestingly, some of the tissue-specific probes (CD235) and all EV-specific probes generated clear signals in fractions 1-5, which are expected to contain the majority of EVs. However, CD63 and CD81 probes, as well as IEL beads for CD171, GAP43 and CD68 detection, all yielded distinctive signals in later fractions (8-17). The localization of L1CAM signal to the late SEC fractions, and the wide distribution of CD63 and CD81, was documented previously (Dmitry Ter-Ovanesyan 2020; Norman et al. 2021) and attributed to the abundance of soluble protein (not specific to EVs). However, such signal could also be generated by an EV subpopulation whose diameter is smaller than anticipated. Our results indicate that at least a large portion of the IEL signal in the late fractions is EV in origin. This notion is supported by evidence. The use of two distinct antigens for particle capture and detection, one of which is a tetraspanin, suggests EV nature. Moreover, electron microscopy revealed vesicles with diameters between 10-30 nm in late fractions, and these fractions contained lipids and mRNA, both characteristic of EVs. Also, pre-processing with detergent prior to IEL analysis reduced the GAP43 positivity in late fraction. The observed shift of CD63 signal from early to late fractions which may be due to the breakdown of larger CD63-positive EVs (originally found in early fractions), followed by re-closure of the membrane fragments to form smaller vesicles. Altogether, the evidence above points to the predominance of small EVs in late fractions, which are detected by IEL. IEL analysis of SEC fractions of the media conditioned by iPSC derived neurons revealed both GAP43 and CD171 signal predominantly in the early (2-5) fractions, and a much weaker signal in the late fractions. This led us to hypothesize that the small EVs are enriched in plasma, either because this fraction has an advantage getting into circulation, or because smaller EVs are generated in the course of penetrating vascular wall (e.g., due to extrusion). The hypothesis that tissue-derived EVs in the plasma are eluted in later fractions is also supported by detection of specific markers of neurons, tissue macrophages and podocytes (not shown) in the late fractions, while the markers of circulating cells, such as erythrocytes, are found in early fractions. Moreover, qPCR analysis showed a much higher ratio between neuronal and oligodendrocyte -specific transcripts, and erythrocyte-specific mRNA in the late fractions, when compared to the early ones. Importantly, TaqMan PCR is an amplicon-based approach and thus further research is needed to determine the presence of full-length and functional mRNA. The main goal of this study was assay development, and further research is needed to reveal the role and biogenesis of the putative novel EV subpopulation that was discovered by the innovative IEL assay described in this work.

Current EV research puts strong emphasis on the heterogeneity of canonical EV markers, such as tetraspanins, in distinct EV populations released from different cell and tissue types. Our novel assay represents a useful new tool to address this problem. We have analyzed the comparative levels of CD63, CD81 and CD9 on the surface of EVs released into plasma by neuronal (GAP43-positive) and erythrocytes (CD235-positive). In agreement with previous findings, erythrocyte specific EVs showed high CD9 content, with relatively low presentation of CD63 and CD81. In contrast, neuronal EVs from both plasma and cell culture presented with high CD63 and CD81 levels, while their CD9 levels were relatively low. These results demonstrate the useful capability of the newly developed IEL for comparative analysis of surface antigen (profiling) of tissue specific EV populations

In conclusion, our novel IEL assay is a sensitive, specific, and user-friendly method for the analysis of EV surface proteins in culture media and especially in biological fluids. It has the advantage of multiplexing, low sample volume requirements and applicability for unprocessed plasma samples. This novel technology enabled the discovery of a smaller subpopulation of tissue specific EVs in blood plasma, which eluted in late SEC fractions, and identified differences in tetraspanin profiles between plasma EV subpopulations. This promising novel technique could result in rapid advancement of EVs research by enabling expedited comprehensive (multi-marker) studies. The assay, which is currently adapted to medium throughput format, could be further expanded towards high-throughput population-wide studies.

## Acknowledgements

TEM samples were prepared and imaged by the Brandeis Electron Microscopy Facility

The study was supported in part by the Intramural Research Program of the National Institute on Aging, NIH (for E. Eren and D. Kapogiannis)

## Declaration of Interest Statement

Olga Volpert, Erez Eitan and Katia Elgart are all employed by NeuroDex Inc., a for-profit start-up company focused on EV-based diagnostic assay development and commercialization, EG was an employee of NeuroDex at the time the work was conducted. EE is also a majority shareholder at NeuroDex.

